# What the book of Lambda doesn’t tell us about temperate phages and lysogeny in the real world

**DOI:** 10.1101/2022.05.11.491505

**Authors:** Brandon A. Berryhill, Rodrigo Garcia, Ingrid C. McCall, Waqas Chaudhry, Marie-Agnès Petit, Bruce R. Levin

## Abstract

The most significant difference between bacteriophages functionally and ecologically is whether they are purely lytic (virulent) or temperate. Virulent phages can only be transmitted horizontally by infection, most commonly with the death of their hosts. Temperate phages can also be transmitted horizontally, but upon infection of susceptible bacteria, their genomes can be incorporated into that of their host’s as a prophage and be transmitted vertically in the course of cell division by their lysogenic hosts. From what we know from studies with the temperate phage Lambda and other temperate phages, in laboratory culture, lysogenic bacteria are protected from killing by the phage coded for by their prophage by immunity; where upon infecting lysogens, the free temperate phage coded by their prophage are lost. Why are lysogens not also resistant as well as immune to the phage coded by their prophage since immunity does not confer protection against virulent phages? To address this question, we used a mathematical model and performed experiments with temperate and virulent mutants of the phage Lambda in laboratory culture. Our models predict and experiments confirm that selection would favor the evolution of resistant as well as immune lysogens, particularly if the environment includes virulent phage that share the same receptors as the temperate. To explore the validity and generality of this prediction, we examined ten lysogenic *Escherichia coli* from natural populations. All ten were capable of forming immune lysogens but their original hosts were resistant to the phage coded by their prophage.

**Significance Statement:** This jointly theoretical and experimental study predicted that lysogenic bacteria, will be resistant as well as immune to the phage coded for by their prophage. All ten naturally occurring lysogenic *Escherichia coli* tested were resistant to the temperate phage coded for by their prophage.

## Introduction

Functionally and ecologically, the single most significant difference among bacteriophages is whether they are virulent (“purely lytic”) or temperate. Virulent phages, a major source of bacterial mortality (1), can only be transmitted horizontally, commonly with the death of the infected bacterium and subsequent release of phage particles (2). Temperate phages can also be transmitted horizontally in this way, but with a low probability, their genomes can become incorporated into that of the infected bacterium as prophage (3), forming lysogens. The infected lysogens survive and the phage genome, borne by these lysogenic bacteria, is transmitted vertically during cell division (3, 4).

Be the phage virulent or temperate, selection will favor bacteria that can survive when confronted by these viruses. Bacteria have several mechanisms that facilitate their survival in the presence of phage. We divide these mechanisms into two functionally distinct classes based on whether or not the phage genome enters the cell. We refer to the mechanisms where the phage genome enters the cells, but the infecting phage are prevented from replicating and are lost, as immunity. Include amongst these immunity mechanisms are CRISPR-Cas (5), restriction-modification (6), abortive infection (7), and super infection exclusion (8). The latter mechanism obtains when lysogens are infected by the phage coded by their prophage. The most prominent mechanism by which phage genomes are prevented from entering cells is receptor site modification, which we define as resistance. Though, other mechanisms for resistance based survival exist such as O-antigen display in which the bacteriophage are sterically inhibited from adsorbing to the bacteria (9). Critical to this definition of resistance is the fact that the phage virion does not attach to the cell and thus the bacteria is refractory to the phage.

Though immunity is the primary mechanism by which lysogens are protected from super infection by the phage coded by their prophage, the vast majority of infections of a naÏve host by a temperate phage result in a lytic cycle. Moreover, virulent mutants of temperate phages, ex-temperate phages (10), can be generated by mutation and there may well be virulent phages that share the same receptor as temperate phages. Why then does selection not also lead to the evolution of resistant lysogens? To address this question, we use a mathematical model of the population and evolutionary dynamics of temperate phages in bacteria capable of generating resistant mutants in sensitive cells and lysogens. The results of our numerical analysis of the properties of our models, simulations, with parameters in the range estimated for *Escherichia coli* and its phages, predict that when populations of sensitive bacteria are confronted by temperate phages, immune lysogens, resistant non-lysogens, and resistant lysogens will emerge and ascend. The ascent of resistant lysogen is particularly prevalent, if the community includes virulent, ex-temperate phages, or other phages that share the same receptors as the temperate. Our tests of this prediction with *E. coli* and temperate and virulent mutants of the phage lambda respectively, Lambda and Lambda^VIR^ are consistent with it.

To explore the real-world generality of this prediction, we examined the susceptibility of ten naturally occurring lysogenic *E. coli* to the phage coded for by their prophage. All ten temperate phages generated by induction of these wild lysogens were capable of infecting and forming immune lysogens in a laboratory strain of *E. coli*. Most critically, as predicted by our models and experiments with the phage Lambda, all ten of the naturally occurring *E. coli* lysogens studied were resistant (refractory) to the phage coded for by their prophage. We discuss the implications of these results to our understanding of the evolution and ecology of lysogenic bacteria and temperate phages.

## Results

### The mathematical model

In Figure 1, we present a diagram of our mathematical model of the population and evolutionary dynamics of virulent and temperate bacteriophages and their bacterial hosts. This model includes populations of non-lysogens (N) and lysogens (L), and free temperate phages (P), as cells and particles per mL, respectively. It also allows for a virulent phage population (V) that shares the same receptor as the temperate phage, and resistant mutants of the lysogens (L_R_), as well as resistant non-lysogens, (N_R_).

**Figure 1.**
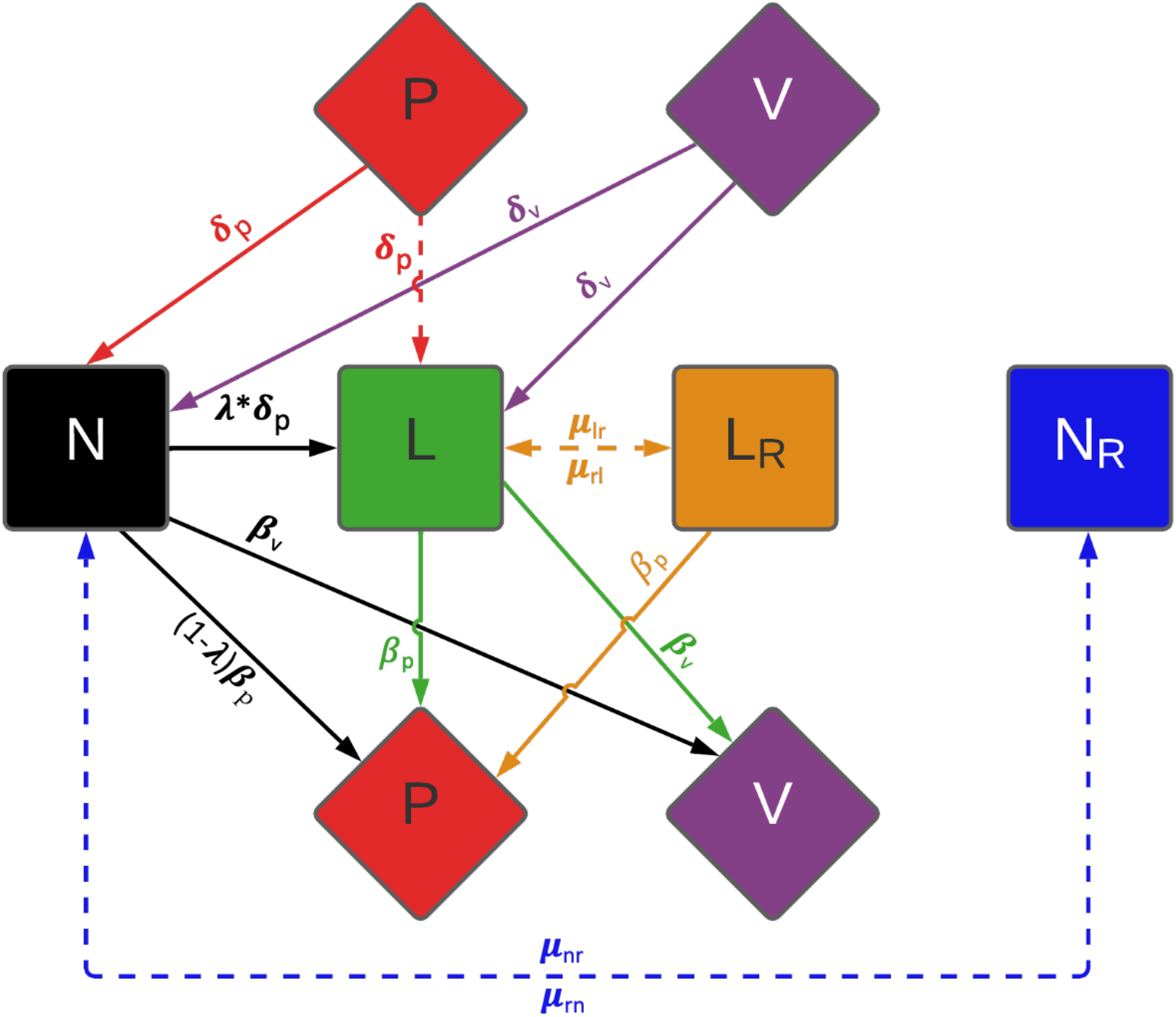
Model of the population and evolutionary dynamics of temperate and lytic phages. There is a population of temperate phage, P; a population of virulent phage, that shares the same receptor with the temperate phage, V; a phage-sensitive non-lysogenic bacterial population, N; a refractory non-lysogenic bacterial population, N_R_; a phage-sensitive lysogenic, L; and a refractory lysogenic bacterial population, L_R_. See the text for a description of the model and Supplemental Table 1 for the definitions, dimensions, of the parameters and the values used in our numerical solutions to the equations, simulations.

Phage infection is a mass action process where the viruses adsorb to the bacteria at rates proportional to the product of the bacterial densities and rate constants δ_p_ and δ_v_, ml attacks per phage particle per cell per hour, for the temperate and virulent phages, respectively (11). With a probability λ Lambda (0 ≤ λ ≤1), infections of sensitive bacteria with temperate phage form lysogens. The remaining fraction (1-λ) of infections by the temperate phages enter a lytic cycle, and each infected bacterium bursts to yield β_P_ phage particles. Infections of sensitive non-lysogenic and lysogenic bacteria with the virulent phages produces β_V_ phage particles.

The rate of growth of the bacteria is equal to the product of their maximum rates v_n_, v_nr_, v_l_, v_lr_ (per hour) and a hyperbolic function, ψ(R), of the concentration of a limiting resource (R, µg/ml) 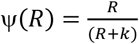, where k is the concentration of the limiting resource where the net growth rate is half its maximum value (12) The limiting resource is consumed at a rate equal to the product of ψ(*R*), the sum of the products of the densities and maximum growth rate of the bacterial populations, and a conversion efficiency parameter, e, µg/cell (13). The rates of mutations L→L_R_,→L_R_→L, N→N_R_, and N_R_→N are, respectively µ_lr_, µ_rl_, µ_nr_ and µ_rn_ per cell per hour (14). To account for the decline in the physiological state as the bacteria approach stationary phase (R = 0), we assume that the rates of phage infection and mutation decline at rates proportional to ψ(R).

With these definitions and assumption, neglecting the lag phase of the bacteria and the latent periods for the phages, and assuming that the likelihood of lysogeny is independent of the multiplicity of infection (15, 16), the rates of change in the concentration of the resource, densities of the different populations of bacteria and the phage are given by the set of coupled differential equations described in supplemental equations sEQ1-sEQ8.

The definitions, dimensions and values of parameters used for our numerical solutions to these equations (simulations) are presented in Supplemental Table 1. Although our primary focus in this study is temperate phages and lysogeny, this is a general model of the population and evolutionary dynamics of bacteria and their virulent and temperate phages. The generality of this model can be seen in Supplemental Figure S1 where we present the predictions of the model and in Figure S2 where we present a set of parallel experiments with *E. coli* and the phages Lambda and Lambda^Vir^ confirming the predictions of the model.

### Model-based predictions and experimental results

We open this consideration with a comparison between the predicted population dynamics generated by numerical solutions to differential equations of this model, simulations, and the corresponding dynamics observed for *E. coli* and the phages Lambda and Lambda^Vir^ in serial transfer culture.

In Figure 2A, we follow the simulated changes in the densities of bacteria and phage in serial transfer population initiated with 10^6^ sensitive bacteria, N, 10^4^ resistant non-lysogens, N_R_, and 10^5^ temperate phage, P. Within short order, lysogens, L, are produced and ascend in density, as do resistant non-lysogens, N_R_. With these parameters and starting conditions, lysogens and resistant non-lysogens become the dominant bacterial populations. A population of resistant lysogens, L_R_ also emerged and increased in frequency, but remained a minority population. As a consequence of induction, free temperate phage are continually produced by the lysogens and resistant lysogens and maintain their population.

**Figure 2.**
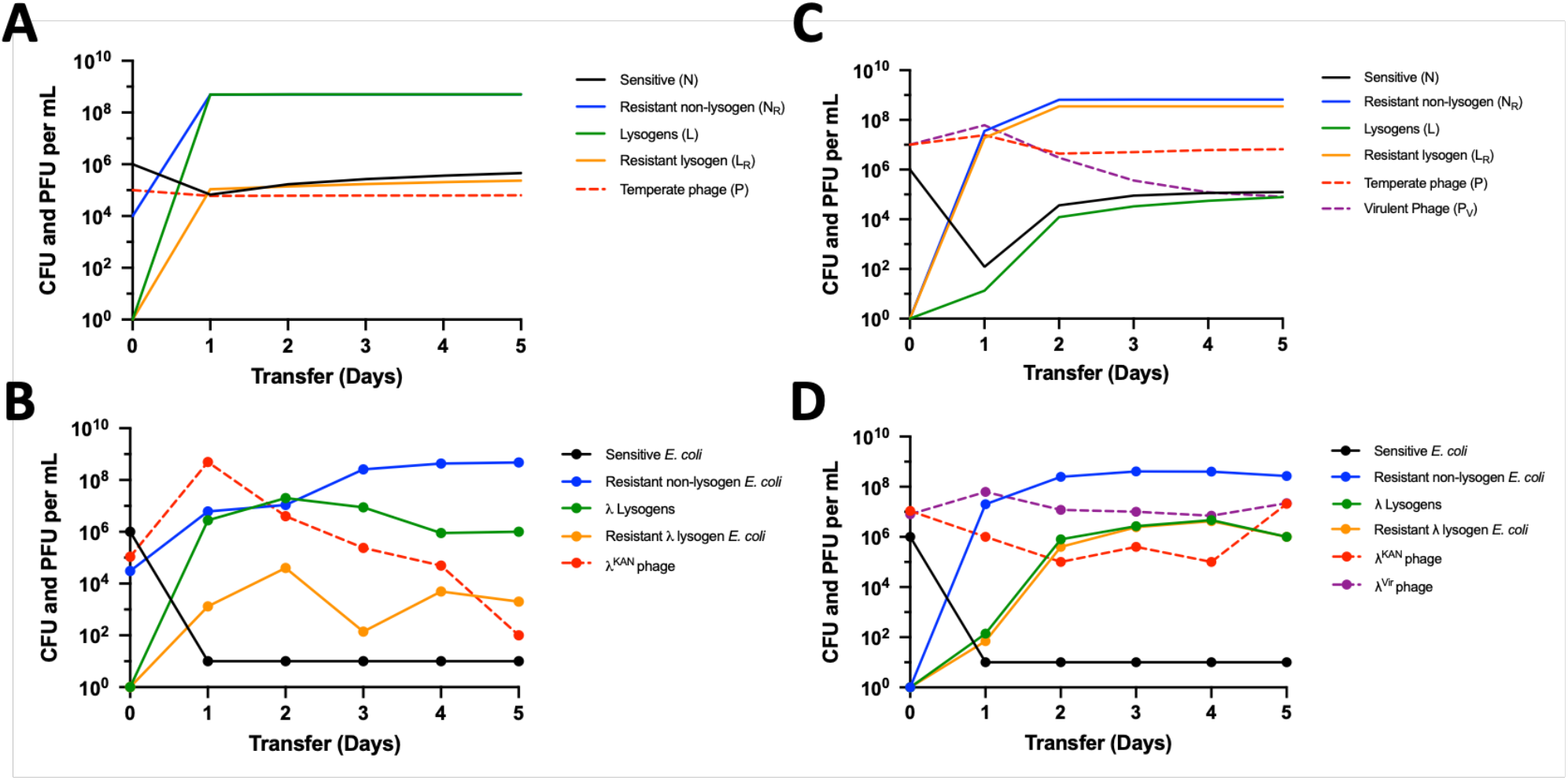
Population dynamics for lysogeny in mixed populations with and without the presence of a virulent phage. (Top section) Simulation results with parameter values in the range estimated for *Escherichia coli* and lambda (Lambda) phage (Table S1). (Lower section) Experiments with sensitive and Lambda-resistant *E. coli*, Lambda^KAN^, and Lambda^Vir^. Changes in bacteria (colony-forming units per mL) and phage (plaque-forming units per mL) densities in a 24 h serial transfer populations with a 1/100 dilution factor are shown for: (**A**) Simulation initiated with 10^6^ sensitive bacteria, and 10^4^ resistant non-lysogens with 10^5^ temperate phage per mL; (**B**) experimental culture initiated with ∼10^6^ Lambda-sensitive *E. coli* and ∼10^4^ Lambda-resistant *E. coli* marked with a streptomycin resistance marker, and ∼10^5^ temperate phage with a kanamycin resistance marker, Lambda^KAN^; (**C**) simulation initiated with 10^6^ phage-sensitive bacteria, 10^7^ temperate phages, and 10^7^ virulent phages that share the same receptor as the temperate phage; (**D**) experimental culture initiated with ∼10^6^ Lambda-sensitive bacteria, ∼10^7^ temperate phage with kanamycin resistance, Lambda^KAN^, and ∼10^7^ virulent mutants of Lambda, Lambda^VIR^.

The parallel serial transfer experiments were initiated with a low density of free Lambda temperate phages, a population of Lambda-sensitive *E. coli*, and a population of Lambda-resistant *E. coli* (Fig. 2B). As predicted by our model, the resistant non-lysogens increased in frequency, as did a novel population that was both resistant (refractory) to phage Lambda and lysogenic for Lambda, resistant lysogens.

To more extensively explore the conditions under which populations of resistant lysogens ascend and are maintained, we initiated our simulated serial transfer populations with sensitive bacteria, temperate phage, and a virulent phage that shares the same receptor as the temperate. Under these conditions, selection for resistance should be more intense. This can be seen in Figure 2C, where resistant lysogens (L_R_), as well as resistant non-lysogens (N_R_), became the dominant bacterial populations. To explore the validity of this prediction experimentally, we initiated serial transfer populations with sensitive *E. coli* and both temperate and virulent Lambda (Fig. 2D). The results of this experiment (and its replicas in Figure S3) are qualitatively consistent with the predictions of the simulations. Resistant non-lysogens, with *malT* mutations similar to what has been reported in other *E. coli* resistant to Lambda-phage ((17, 18); Fig. S4), ascend and become the dominant bacterial population followed by lysogens and resistant lysogens (see Fig. S5 for the evidence that the resistant lysogens appear refractory).

Our model and experiments make two predictions: (i) When a population of sensitive bacteria are infected by temperate phages, in addition to the generation and ascent of lysogens, resistant lysogens will be generated and increase in density; (ii) If the community includes virulent phages that share the same receptor as the temperate phage, the resistant lysogens will ascend and become the dominant population of lysogenic bacteria. To explore the validity and generality of these predictions, we used a set of naturally occurring (wild) lysogenic *E. coli* with different prophages from sewage and the gut microbiomes of infants (19) (Table S2). We induced these wild lysogens to produce free temperate phages which we then used to infect the lysogens from whence they came.

Consistent with the prediction of our model and experiments with Lambda, all ten naturally occurring lysogens were refractory to the free phages generated by induction of their host lysogens (Figure 3A). To determine whether this resistance is property of the host bacteria rather than the prophage, Lambda-sensitive *E. coli* C, Lambda lysogens, and Lambda^Vir^-resistant bacteria were infected with a temperate Lambda-phage coding for kanamycin resistance, Lambda^KAN^. As anticipated, the phages replicated on the sensitive non-lysogens but not the Lambda^Vir^-resistant *E. coli*. However, not anticipated was the marked increase in the density of free phages in a control experiment with Lambda lysogens infected by temperate Lambda-phage (Figs. 3A and 3B, shaded areas).

**Figure 3.**
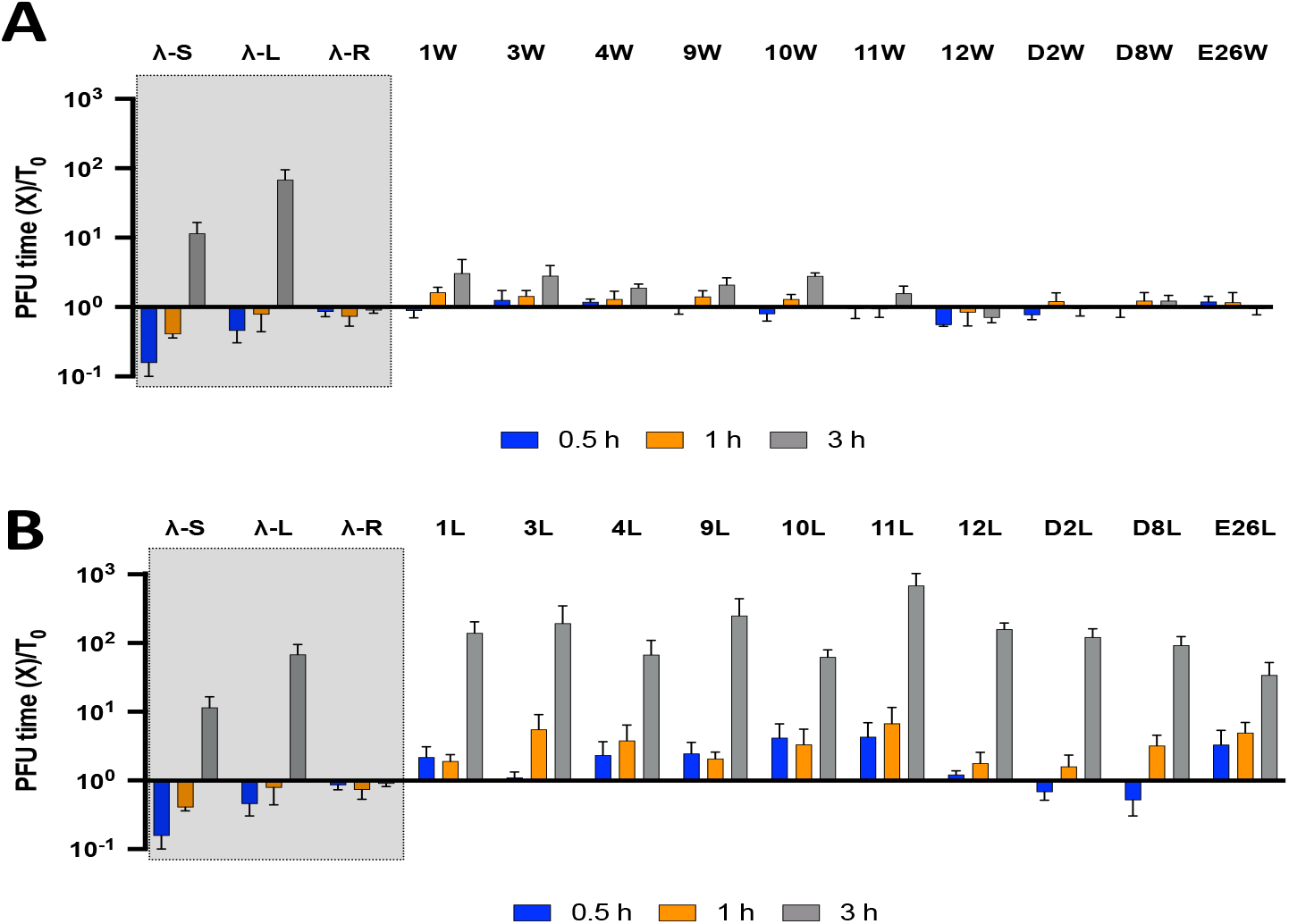
Susceptibility and resistance of naturally occurring and *E. coli* C constructed lysogens. Ratios of free phage density after 0.5, 1, and 3 h relative to that at 0 h (T_0_). Means and standard deviations of the ratios of three replicas of this experiment. Shaded regions infections with phage lambda (Lambda) in three distinct bacterial states: Lambda-sensitive (Lambda-S), Lambda-lysogen (Lambda-L), and Lambda-resistant (Lambda-R). (A) Naturally occurring lysogens (W) infected with a low multiplicity of infection of the induced free phages for which they are lysogenic. (B) *E. coli* C lysogens constructed (L) from the prophage induced from the corresponding wild lysogen (W) infected with a low multiplicity of induced wild phages for which they are lysogenic. See Table S1 for designations and sources of the naturally occurring lysogenic *E. coli*.

To test the hypothesis that the resistance of naturally occurring lysogens to the phages coded by their prophages is a property of the bacteria rather than that of the prophage, we infected *E. coli* C with the temperate phages induced from the naturally occurring lysogens. None of the ten wild prophage bearing lysogens of *E. coli* C, were resistant (refractory) to the free phages coded by their prophage (Fig. 3B). They behaved like Lambda lysogens infected with Lambda (Fig. 3B, shaded area). Most intriguingly, similar to the Lambda lysogens in the control experiment (shaded areas of Fig. 3), when the *E. coli* C wild phage lysogens were infected with a low density of free temperate phages, high densities of free phages were produced.

The observation that when infected with free phages both the Lambda lysogens, and the *E. coli* C lysogens bearing the wild temperate phage produce high densities of free phages (Fig. 3B) was unexpected. It suggested that lysogens were induced when infected with the temperate phages coded by their prophage. We postulate that infections with temperate phages induce lysogens by mounting an SOS response, as would be the case when lysogens are exposed to ultraviolet light or other SOS-inducing insults (20-22). To test this hypothesis, a Δ*recA E. coli* construct obtained from the KEIO collection was lysogenized with Lambda and reinfected with free Lambda phage (23). Contrary to that observed with the *recA*^+^ lysogen (Fig. 3B), infections of the Δ*recA* Lambda lysogen with free Lambda-phage did not generate a high density of free phage (Fig. S6A). To further test the hypothesis that infections with free Lambda induce Lambda lysogens by generating an SOS response, we transformed the Δ*recA E. coli* Lambda lysogens with plasmids either lacking or bearing *recA* (Fig. S7). With the cells bearing a functional *recA* in *trans*, the lysogens were induced at a high rate (Fig. S6B). This did not occur with the plasmid without *recA* (Fig. S6C).

## Discussion

Bacteria are protected from being killed by infecting phage via two distinct mechanisms: immunity and resistance. In the case of immunity, the phage injects its genetic material, but phage replication does not occur, and the infecting phage are lost. This is the case for defense systems such as CRISPR-Cas (5), restriction modification (6), abortive infection (7), and lysogeny (8). In the case of resistance, the phage virion fails to bind to the host bacterium, for example due to modification of the receptor site in the bacteria or the display of the O-antigen (9). For resistance, the bacteria are refractory to the bacteriophage and the phage are not lost.

When sensitive populations of bacteria are confronted with virulent phages, selection favors the evolution of mutants that are resistant to those phages or immune. In accord with what is known about lysogeny, what “the book of Lambda” tells is, when populations of sensitive bacteria are confronted with temperate phage, a fraction of the infected bacteria acquire and maintain the genome of the phage, as a prophage, and the resulting lysogens are immune to super-infection with the phage coded for by their prophage; the phage infecting lysogens are lost. What “the book of Lambda” does not say is that when populations of sensitive bacteria are exposed to temperate phages, resistant bacteria will also emerge and ascend, as presented in this theoretical and experimental study with *E. coli* and Lambda phage. Moreover, if the community contains virulent phages that share the same receptors as the temperate phages coded by the prophage of lysogens, resistant lysogens will emerge and rise to become the dominant population of lysogens.

To explore the generality of this prediction in the real world, by induction we isolated temperate phage from ten naturally occurring lysogens. All ten of these “wild” temperate phages were capable of forming immune lysogens with a laboratory strain of *E. coli*. And, as predicted by our models and experiments with *E. coli* and Lambda, all ten of these naturally occurring lysogens were resistant (refractory) to the phage coded for by their prophage. Based on these theoretical and experimental results, we predict that naturally occurring lysogenic bacteria will be resistant as well as immune to the phage coded for by their prophage.

The results of this study provide another observation not anticipated from the “book of Lambda”. It is known that the rate of induction of prophage is augmented when lysogens are infected with phages other than those coded by their prophage (24, 25). The results of our assay indicated that the rate of induction of lysogens is also increased due to phage infection with the temperate phages coded by their prophage. Our analysis indicates that by promoting an SOS response, infections of lysogens by free phage increase the rate of induction and the production of free phage by populations of lysogens.

The subtitle of a not-quite-ancient theoretical study of the population dynamics of lysogeny raised the question of “why be temperate?” (13). In retrospect, it would seem the interesting question that should have been raised is “why be virulent”? From an ecological and evolutionary perspective, a temperate mode of replication and transmission would be better than a virulent one for a phage. Unless their rates of adsorption and burst sizes are markedly less than that of virulent phage, when invading a population of sensitive bacteria temperate phage will be spread nearly as fast as virulent. By forming lysogens and thereby being transmitted vertically, temperate phages can be maintained in the community when the density of their host population is too low for them to persist as virulent phage (11, 26). Moreover, the phage genome can also increase in frequency when the prophage encodes a phenotype that augments the fitness of its host bacterium (27). Resistance to the phages coded for by their prophage, would be an additional asset to temperate phages. The prophage genome will be transmitted vertically in the course of cell division and the free temperate phages will be continually produced by induction. These free temperate phages will then be capable of infecting sensitive non-lysogens, thereby expanding the range of bacteria bearing that prophage. These lysogens, and thereby the temperate phages they encode, will not be subject to extinction by virulent phages capable of adsorbing to the same receptors as those of the temperate phages.

The molecular biologist Jacques Monod quipped that “What is true for *E. coli* is true for elephants, only more so” (28). While we appreciate this form of inductive inference, we understand that all of the experiments reported here were conducted only with *E. coli*. We postulate that naturally occurring lysogenic bacteria of any species that can produce resistant mutants will be resistant to the phages coded by their prophage.

## Materials and Methods

### Numerical solutions (simulations)

For our numerical analysis of the properties the model of population and evolutionary dynamics of temperate and virulent phage (Figure 1 and equations 1-7), were solved with Berkeley Madonna, using parameters in the ranges estimated experimentally with *E. coli* and virulent and temperate Lambda phages (Table 1). In these numerical solutions (simulations), the changes in states, N→N_R_, N_R_→N, L→L_R_ and L_R_→L are stochastic processes modeled with a Monte Carlo protocol. All other changes are deterministic. Copies of the Berkeley Madonna program used for these simulations are available at www.eclf.net.

### Growth media

LB broth (244620) was obtained from BD. The DM (Davis Minimal) minimal base without dextrose (15758-500G-F) was obtained from Sigma Aldrich. LB agar (244510) for plates was obtained from BD. Phage plates are prepared with 1g/L Yeast Extract (BD-241750), 10 g/L Bacto-Tryptone (Fischer-BP1421), 8 g/L NaCl (Fischer-S271), 10 g/L Agar (BD-214010), 1g/L Glucose (Sigma-G5767) and 2 mM CaCl_2_ (Sigma-C5080). Double-layer soft agar was prepared with 1 g/L Yeast Extract, 10 g/L Bacto-Tryptone, 8 g/L NaCl, 7 g/L Agar, 1g/L Glucose and 2 mM CaCl_2_.

### Bacterial strains

*E. coli* MG1655 was obtained from the Levin Lab Bacterial collection. The wild strains and *E. coli* C were obtained from Marie-Agnès Petit. All lysogens were made in the Levin Lab and saved at −80°C. *E. coli* ΔRecA was obtained from the Keio collection.

### Phages

Temperate Lambda, Lambda^KAN^, and Lambda^VIR^ were obtained from Maroš Pleška at The Rockefeller University.

### Plasmids

All inserts were synthesized and cloned into the plasmids by GenScript (NJ, USA).

### Primers

All primers were ordered from Integrated DNA Technologies (Iowa, USA) as custom oligos with standard desalting.

### Antibiotics

Ampicillin (A9518), Kanamycin (127K0058) and Streptomycin (081K1275) were obtained from Sigma Aldrich (USA).

### Sampling bacterial and phage densities

The densities of bacteria and phage were estimated by serial dilution in 0.85% saline and plating. The total density of bacteria was estimated on Lysogeny Broth (LB) IN hard (1.6%) agar plates. In competition experiments with multiple bacterial populations, diluted samples were plated on LB hard (1.6%) agar plates supplemented with kanamycin (2.5 ug/mL) or streptomycin (4 ug/mL) to distinguish between sensitive, resistant, and lysogenic bacteria. To estimate the densities of free phage, chloroform was added to suspensions before serial dilution. These suspensions were plated at various dilutions on lawns with 0.1 mL of overnight Davis Minimal (DM) Glucose-grown cultures (about 5×10^8^) of *E. coli* MG1655 or *E. coli* MG1655 lysogenized with lambda (to distinguish Lambda^Vir^ from temperate Lambda) and 4mL of LB soft (0.7%) agar on top of hard (1.6%) LB agar plates.

### Testing for Resistance

Bacteria were tested for resistance to Lambda^VIR^ by streaking colonies across a line made with 200 μL of a Lambda^Vir^ lysate (>10^8^ pfu/mL) on LB hard (1.6%) agar plates. Continuous lines were interpreted as evidence for resistance and broken lines sensitivity.

### Testing for whether the bacteria are refractory or immune to the phage

High densities of bacteria were exposed to phage at a low multiplicity of infection (0.01-0.1) in 10ml of DM-Glucose media. The density of phage in these cultures were estimate at the time of first exposures, T=0, and T= 0.5, 1.0 and 3.0 hours later. The bacteria were considered refractory to the phage if there was little or no net change in their density. The bacteria were considered immune, if the density of phage declined or increased in later samples.

### Growth rate estimation

Exponential growth rates were estimated from changes in optical density (OD_600_) in a Bioscreen C. For this, 48-hour stationary phase cultures were diluted in DM glucose liquid media to an initial density of approximately 10^5^ cells per mL. Ten replicas were made for each estimate by adding 300µl of the suspensions to the wells of the Bioscreen plates. The plates were incubated at 37°C and shaken continuously. Estimates of the OD (600nm) were made every five minutes.

### Burst size estimation

The burst sizes for Lambda and Lambda^Vir^, β_p_ and β_v_, were estimated with a one-step growth protocol similar to that first presented in 1939 (Ellis and Delbrück, 1939) and employed in (Stewart et al. 1977). Exponentially growing cultures of *E. coli* MG1655 in DM minimal glucose were used as the host bacteria for these estimates.

### Serial transfer experiments

All serial transfer experiments were carried out in 50 ml Erlenmyer flasks with 10 mL DM minimal glucose media at 37 °C with vigorous shaking. Unless otherwise noted, these serial transfer cultures were initiated by a 1:100 dilution from 10-mL overnight cultures grown from single colonies. Phages were added to these cultures at the initial densities shown in the figures. At the end of each transfer, 0.1 mL of the cultures were transferred into flasks with fresh medium (1:100 dilution). Subsequently, 0.1 mL samples were taken to estimate the densities of bacteria and phage.

### Sequencing and analyses

Phage DNA was extracted using Invitrogen’s (California, USA) PureLink Viral RNA/DNA extraction kit (Cat# 12280-050) using the manufacturer’s protocol and bacterial DNA was extracted using Promega’s (Wisconsin, USA) Wizard Genomics DNA Purification Kit (Cat# A1120) using the manufacturer’s protocol. The extracted DNA was quantified on ThermoFisher’s NanoDrop One^C^ microvolume spectrophotometer (Cat# ND-ONE-W). Samples were sent to the Microbial Genome Sequencing Center in Pittsburgh, Pennsylvania, USA, for whole genome sequencing on the Illumina NextSeq 2000 platform. Analysis of FASTAq files received from the Microbial Genome Sequencing Center were analyzed using Geneious Prime version 2022.0.1.

### Primer design and PCR

Primers were designed using PrimerBLAST (NCBI). PCR was carried out using Thermo Scientific’s (Lithuania) Phusion Blood Direct PCR Master Mix (Cat# F-175L). Products were visualized on a 1% agarose/TAE gel with Biotium’s (California, USA) 10,000X GelRed Nucleic Acid Stain.

### RecA-expressing plasmid construction

A pMAL-c4X-RecA plasmid was synthetized by GenScript (NJ, USA) using as a backbone the plasmid pMAL-c4X (Addgene #75288) and cloning between the NdeI and BamHI restriction sites in the plasmid a *recA*-tt construct (1,196 bp) designed with the *recA* CDS (1,059 bp, Genbank: EU896799.1), a TAGTAGAG linker sequence (BioBrick standard RFC[10]) added to the 3’-terminal end of the *recA* CDS stop codon followed by a downstream double-terminator sequence (BioBrick Part:BBa_B0015). This puts the *recA* CDS under the control of the lactose-inducible TAC promoter, therefore RecA can be complemented in *trans* with induction by IPTG.

### RecA complementation assay

Plasmids pMAL-c4X-RecA and pMAL-c4X were electroporated with 1 pulse of 2.5 kV using a MicroPulse electroporator (Bio-rad, USA) into electrocompetent Lambda-lysogenized *E. coli* Δ*recA* cells prepared from an overnight growth in LB media, followed by three cycles of centrifugation at 13,000g and a wash with ice-cold 10% glycerol solution. Transformants were selected on LB plates supplemented with 12 ug/mL Ampicillin and incubated at 37ºC overnight, colonies were then streaked two times for isolation.

To confirm the presence of the plasmid, Colony-PCR (described previously) was performed using the M13/pUC_Reverse (5’-AGCGGATAACAATTTCACACAGG-3’) and M13/pUC_Forward (5’-CCCAGTCACGACGTTGTAAAACG-3’) primers. The IPTG-induction of the *recA* gene in the complemented Lambda-lysogenized strain *E. coli* Δ*recA* / pMAL-c4X-RecA was performed in a culture of 100 mL LB broth supplemented with 15 ug/mL Ampicillin inoculated with a 5% volume from an overnight culture, the cells were grown aerobically at 37ºC until an OD_600_ of 0.3 and growth was maintained for 3 hours after the addition of 10 mM Isopropyl β-D-1-thiogalactopyranoside (IPTG). The refractory phenotype (described previously) was confirmed directly from IPTG-induced Lambda-lysogenized cultures, using as plasmid-free and insert-free controls the strains *E. coli* Δ*recA*, and *E. coli* Δ*recA* / pMal-c4X, respectively.

### Resistance genotype analysis

The 2,706 bp region of the *malT* gene of the sensitive non-lysogens (MalT WT) and resistant non-lysogens (MalT R) were amplified via two-overlapping PCRs with primers shared by Marie-Agnès Petit. The first product using the primers malT up (5’-GTGACACAGTGCAAATTCAG-3’) and malTint down (5’-CTAGCAGGGTGTTAACTTC-3’) amplifies a 1,297 bp 5’-end region of the *malT* gene with a 138 bp upstream non-coding segment, the second product used the primers Int-up malt (5’-TCCGCAGTTGGTGTTATTG-3’) and malTdown2 (5’-GGTGCGGTTTAGTTTGATAG-3’) to amplify an 1,483 bp 3’-end region of the *malT* gene and a 97 bp downstream segment. These products were purified by running on a 1% agarose gel and gel extracted using the GenElute™Gel Extraction kit (Sigma-Aldrich, USA). The extracted bands were Sanger sequenced by Eurofins (USA) and the resulting sequences analyzed, and a full *malT* gene sequence is obtained with the pairwise alignment of both products. The *malT* sequences were *in silico* translated with an *E. coli* translation table to obtain the amino acid sequences of the MalT receptors, alignments were generated using Clustal Omega 1.2.3 using default parameters and a graphical representation of the alignment was obtained using Geneious 2022.0.1

### Growth media

LB broth (244620) was obtained from BD. The DM (Davis Minimal) minimal base without dextrose (15758-500G-F) was obtained from Sigma Aldrich. LB agar (244510) for plates was obtained from BD. Phage plates are prepared with 1g/L Yeast Extract (BD-241750), 10 g/L Bacto-Tryptone (Fischer-BP1421), 8 g/L NaCl (Fischer-S271), 10 g/L Agar (BD-214010), 1g/L Glucose (Sigma-G5767) and 2 mM CaCl_2_ (Sigma-C5080). Double-layer soft agar was prepared with 1 g/L Yeast Extract, 10 g/L Bacto-Tryptone, 8 g/L NaCl, 7 g/L Agar, 1g/L Glucose and 2 mM CaCl_2_.

### Bacterial strains

*E. coli* MG1655 was obtained from the Levin Lab Bacterial collection. The wild strains and *E. coli* C were obtained from Marie-Agnès Petit. All lysogens were made in the Levin Lab and saved at −80°C. *E. coli* ΔRecA was obtained from the Keio collection.

### Phages

Temperate Lambda, Lambda^KAN^, and Lambda^VIR^ were obtained from Maroš Pleška at The Rockefeller University.

### Plasmids

All inserts were synthesized and cloned into the plasmids by GenScript (NJ, USA).

### Primers

All primers were ordered from Integrated DNA Technologies (Iowa, USA) as custom oligos with standard desalting.

### Antibiotics

Ampicillin (A9518), Kanamycin (127K0058) and Streptomycin (081K1275) were obtained from Sigma Aldrich (USA).

## Acknowledgments

The authors thank Melony Ivey, Esther Lee, Adithi Govindan, Nicole Vega, Ross Greenberg, Joanna Goldberg, David Goldberg, Thomas O’Rourke, Teresa Gil-Gil, Eduardo Rodriguez-Roman, and Andrew Smith for their comments on the manuscript, feedback on experiments, and support in the laboratory.

## Data and materials availability

The Berkeley Madonna program used for the simulations are available at www.eclf.net. The sequence data shown have been deposited at Genbank (NCBI, Bethesda, Maryland, USA). All other data are available in the main text or supplementary materials.

