## Supplemental Materials for "What the book of Lambda doesn’t tell us about temperate phages and lysogeny in the real world"

**This PDF file includes:**

Supplementary equations
Figures S1 to S7 (not allowed for Brief Reports)
Tables S1 to S2 (not allowed for Brief Reports)
SI References

**Supplemental Equations**

$\boldsymbol{\Psi(R)} = \frac{\boldsymbol{R}}{\boldsymbol{K+R}}$ sEQ. 1

$\frac{dR}{dt} = -\Psi(R) \cdot e \cdot (v \cdot N + v_{nr} \cdot N_R + v_l \cdot L + v_{lr} \cdot L_R)$ sEQ. 2

$\frac{dN}{dt} = \Psi(R) \cdot \left(v \cdot N - \delta_v \cdot N \cdot V - \delta_p \cdot N \cdot P - \mu_{nr} \cdot N + \mu_{rn} \cdot N_R\right)$ sEQ. 3

$\frac{dN_R}{dt} = \Psi(R) \cdot (v_{nr} \cdot N_R + \mu_{nr} \cdot N - \mu_{rn} \cdot N_R)$ sEQ. 4

$\frac{dL}{dt} = \Psi(R) \cdot (v_l \cdot L + \lambda \cdot \delta_p \cdot N \cdot P - \delta_v \cdot V \cdot L - \gamma \cdot L - \mu_{lr} \cdot L + \mu_{rl} \cdot L_R$ sEQ. 5

$\frac{dL_R}{dt} = \Psi(R) \cdot (v_{lr} \cdot L_R - \gamma \cdot L_R - \mu_{rl} \cdot L_R + \mu_{lr} \cdot L)$ sEQ. 6

$\frac{dP}{dt} = \Psi(R) \cdot \left(\delta_p \cdot N \cdot (1 - \lambda) \cdot P \cdot \left(\beta_p - 1\right) - \delta_p \cdot L \cdot P + \gamma \cdot (L + L_R) \cdot \beta_p\right)$ sEQ. 7

$\frac{dV}{dt} = \Psi(R) \cdot \delta_v \cdot (N + L) \cdot (\beta_v - 1) \cdot V$ sEQ. 8

### Supplemental Figures

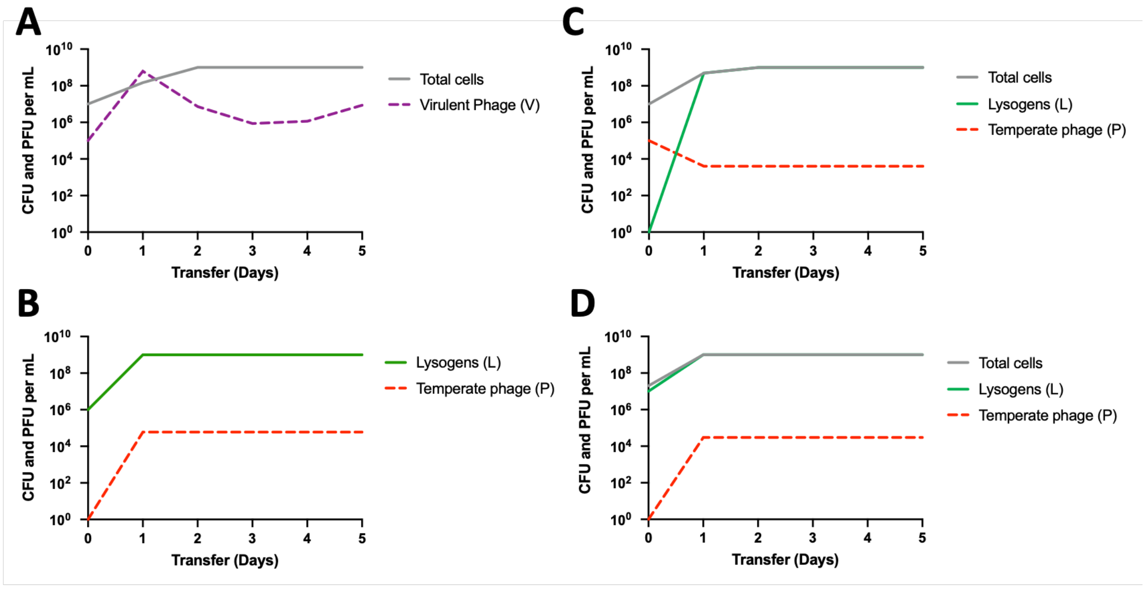


**Fig. S1. Simulation results for basic phage-bacteria interactions.** Shown are the simulated changes in the densities of phage (PFU/mL) and bacteria (CFU/mL) in serial transfer culture. Save for time zero, the densities are presented as the output at each 24-hour interval. (A) A simulation of $10^7$ sensitive cells being confronted with $10^5$ virulent phages (dashed purple). Shown are the density of phages and the total density (grey) of cells ($N + N_r$). (B) A simulation initiated with $10^6$ lysogenic bacteria (green) with free temperate phages (dashed red) being produced by spontaneous induction. (C) A simulated population initiated with $10^7$ sensitive cells being confronted by $10^5$ temperate phages. Shown are the densities of lysogenic bacteria (green), temperate phage (dashed red), and total cells ($N + L + N_R$, in grey). (D) A simulation started with equal densities ($10^7$) of sensitive cells and lysogens. Shown are the lysogens (green) and temperate phage (dashed red) populations as well as total cells ($N + L$, in grey).

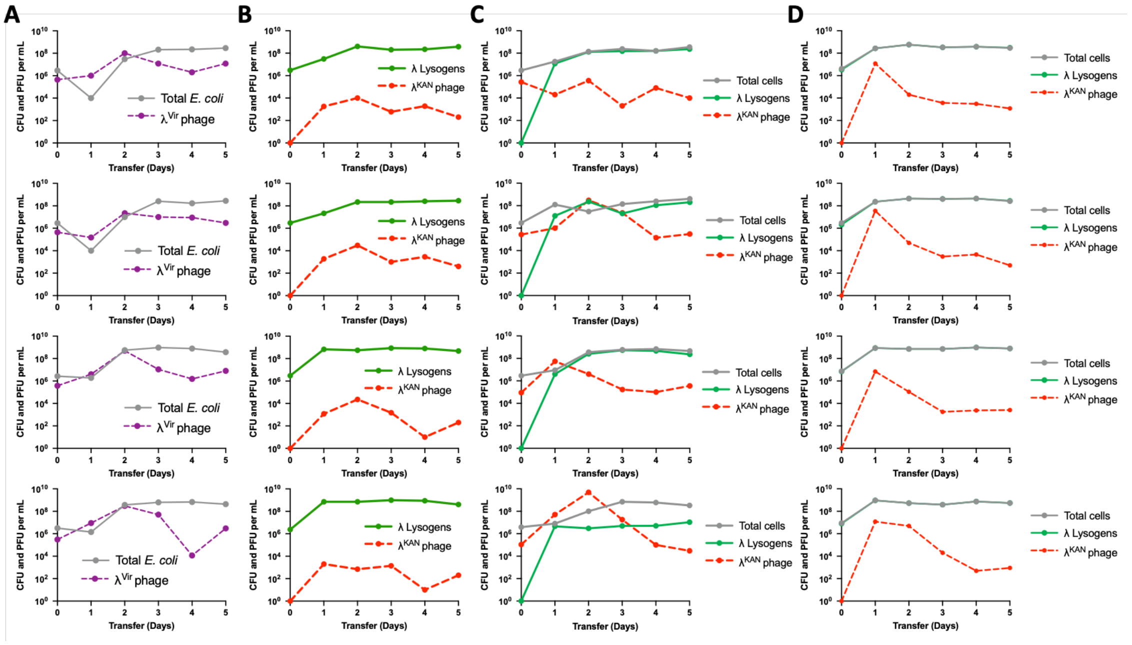


**Fig. S2. Experimental exploration of the predictions of the model.** Presented are four biological replicas of four experiments in parallel to the modelling results in Figure 2. Shown are the changes in the densities of phage (PFU/mL) and bacteria (CFU/mL) in serial transfer culture sampled at 24-hour intervals. (A) Cultures of ~$10^7$ sensitive *E. coli* confronted with ~$10^5$ virulent phages. Shown are the densities of virulent phage (dashed purple) and the total cell population (grey). (B) A population of ~$10^6$ sensitive λ lysogens (green) growing in the absence of phage and generating free temperate $\lambda^{KAN}$ phages (dashed red) via spontaneous induction. (C) A population of ~$10^7$ sensitive *E. coli* confronted with $10^5$ $\lambda^{KAN}$ phages. Shown are the densities of total cells (grey), *E. coli* lysogenic for $\lambda^{KAN}$ (green), and the density of free $\lambda^{KAN}$ phage (dashed red). (D) Experimental cultures started with equal densities ($10^7$) of sensitive non-lysogenic *E. coli* and sensitive $\lambda^{KAN}$ lysogens. Shown are the lysogens (green) and temperate phage (dashed red) populations as well as total cells (grey). Total cells are superimposed on top of lysogenic *E. coli*.

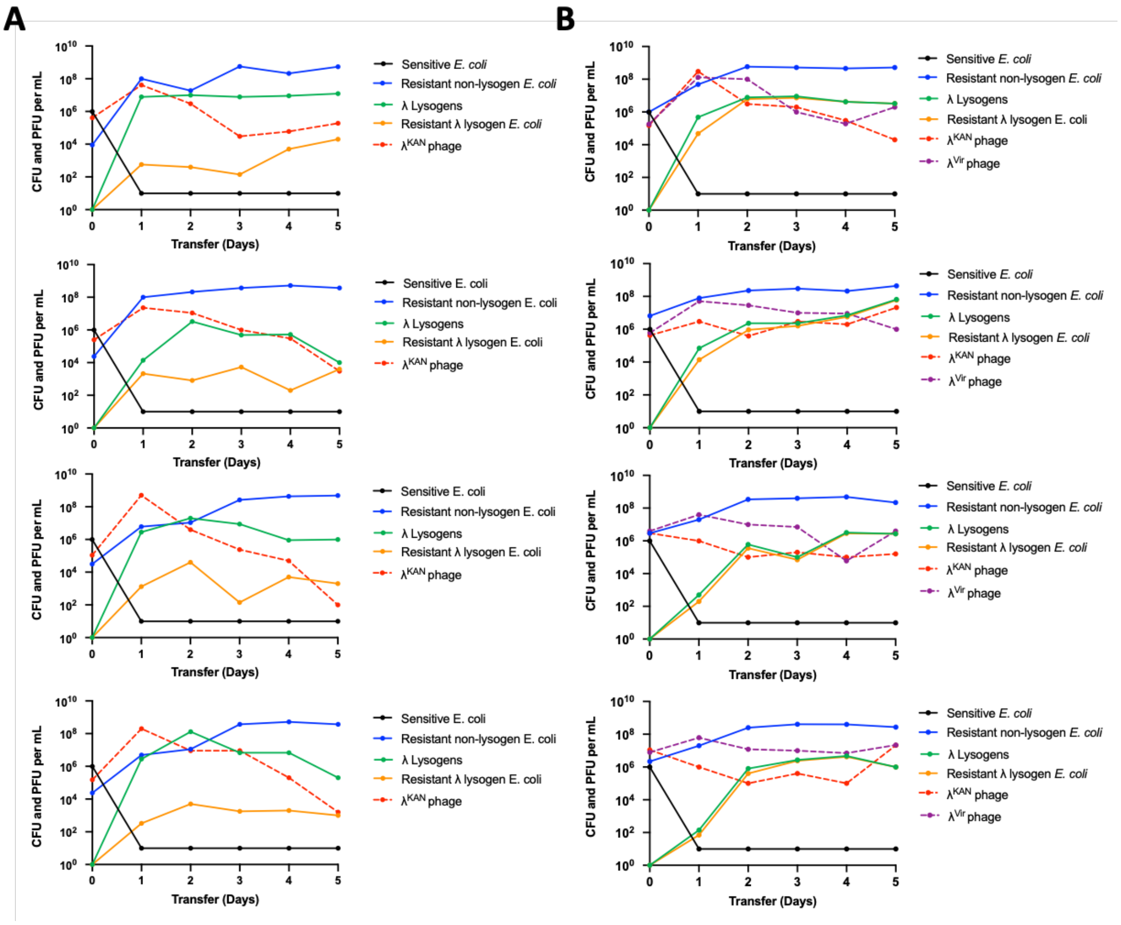


**Fig. S3. Replicas of Figures 2B and 2D.** Four biological replicas of experiments initiated with sensitive (black) and resistant non-lysogenic *E. coli* (blue), $\lambda^{KAN}$ (dashed red), and $\lambda^{Vir}$ (dashed purple), notable is the emergence of the resistant λ lysogenic *E. coli* (orange) . Changes in the densities of bacteria (CFU per mL) and phage (PFU per mL) in 24-hour serial transfer populations with a 1/100 dilution factor are shown for: (A) Experimental cultures initiated with ~$10^6$ λ-sensitive *E. coli*, and ~$10^4$ lambda-resistant $STR^R$ *E. coli*, and ~$10^5$ temperate phage coding for kanamycin resistance, $\lambda^{KAN}$. (B) Experimental culture initiated with ~$10^6$ λ-sensitive bacteria, ~$10^7$ temperate phage coding for kanamycin resistance temperate phage, $\lambda^{KAN}$ and ~$10^7$ virulent mutants of lambda, $\lambda^{Vir}$ .

```
                   10         20         30         40         50       60
Consensus MLIPSKLSRP VRLDHTVVRE RLLAKLSGAN NFRLALITSP AGYGKTTLIS QWAAGKNDIG
MalT WT   MLIPSKLSRP VRLDHTVVRE RLLAKLSGAN NFRLALITSP AGYGKTTLIS QWAAGKNDIG
MalT R    MLIPSKLSRP VRLDHTVVRE RLLAKLSGAN NFRLALITSP AGYGKTTLIS QWAAGKNDIG

                   70         80         90        100        110        120
Consensus WYSLDEGDNQ QERFASYLIA AVQQATNGHC AICETMAQKR QYASLTSLFA QLFIELAEWH
MalT WT   WYSLDEGDNQ QERFASYLIA AVQQATNGHC AICETMAQKR QYASLTSLFA QLFIELAEWH
MalT R    WYSLDEGDNQ QERFASYLIA AVQQATNGHC AICETMAQKR QYASLTSLFA QLFIELAEWH

                  130        140        150        160        170        180
Consensus SPLYLVIDDY HLITNPVIHE SMRFFIRHQP ENLTLVVLSR NLPQLGIANL RVRDQLLEIG
MalT WT   SPLYLVIDDY HLITNPVIHE SMRFFIRHQP ENLTLVVLSR NLPQLGIANL RVRDQLLEIG
MalT R    SPLYLVIDDY HLITNPVIHE SMRFFIRHQP ENLTLVVLSR NLPQLGIANL RVRDQLLEIG

                  190        200        210        220        230        240
Consensus SQQLAFTHQE AKQFFDCRLS SPIEAAESSR ICDDVSGWAT ALQLIALSAR QNTHSAHKSA
MalT WT   SQQLAFTHQE AKQFFDCRLS SPIEAAESSR ICDDVSGWAT ALQLIALSAR QNTHSAHKSA
MalT R    SQQLAFTHQE AKQFFDCRLS SPIEAAESSR ICDDVSGWAT ALQLIALSAR QNTHSAHKSA

                  250        260        270        280        290        300
Consensus RRLAGINASH LSDYLVDEVL DNVDLATRHF LLKSAILRSM NDALITRVTG EENGQMRLEE
MalT WT   RRLAGINASH LSDYLVDEVL DNVDLATRHF LLKSAILRSM NDALITRVTG EENGQMRLEE
MalT R    RRLAGINASH LSDYLVDEVL DNVDLATRHF LLKSAILRSM NDALITRVTG EENGQMRLEE

                  310        320        330        340        350        360
Consensus IERQGLFLQR MDDTGEWFCY HPLFGNFLRQ RCQWELAAEL PXXXXXXAXX XXXXXXXXXX
MalT WT   IERQGLFLQR MDDTGEWFCY HPLFGNFLRQ RCQWELAAEL PEIHRAAAES WMA---QGFP
MalT R    IERQGLFLQR MDDTGEWFCY HPLFGNFLRQ RCQWELAAEL PVGTGGGAAG NPPCRRRKLD

                  370        380        390        400        410        420
Consensus XXXIXXXXXX XXAGXXXXXX XXXXXXXXSX XNHSELSLLE ESLKALPWDS LLENPQLVLL
MalT WT   SEAIH---HA LAAGDALMLR DILLNHAWSL FNHSELSLLE ESLKALPWDS LLENPQLVLL
MalT R    GPGISQRSNS SCAGGRRCAD A----ARYSA *--------- ---------- ----------

                  430        440        450        460        470        480
Consensus QAWLMQSQHR YGEVNTLLAR AEHEIKDIRE DTMHAEFNAL RAQVAINDGN PDEAERLAKL
MalT WT   QAWLMQSQHR YGEVNTLLAR AEHEIKDIRE DTMHAEFNAL RAQVAINDGN PDEAERLAKL
MalT R    ---------- ---------- ---------- ---------- ---------- ----------

                  490        500        510        520        530        540
Consensus ALEELPPGWF YSRIVATSVL GEVLHCKGEL TRSLALMQQT EQMARQHDVW HYALWSLIQQ
MalT WT   ALEELPPGWF YSRIVATSVL GEVLHCKGEL TRSLALMQQT EQMARQHDVW HYALWSLIQQ
MalT R    ---------- ---------- ---------- ---------- ---------- ----------

                  550        560        570        580        590        600
Consensus SEILFAQGFL QTAWETQEKA FQLINEQHLE QLPMHEFLVR IRAQLLWAWA RLDEAEASAR
MalT WT   SEILFAQGFL QTAWETQEKA FQLINEQHLE QLPMHEFLVR IRAQLLWAWA RLDEAEASAR
MalT R    ---------- ---------- ---------- ---------- ---------- ----------

                  610        620        630        640        650        660
Consensus SGIEVLSSYQ PQQQLQCLAM LIQCSLARGD LDNARSQLNR LENLLGNGKY HSDWISNANK
MalT WT   SGIEVLSSYQ PQQQLQCLAM LIQCSLARGD LDNARSQLNR LENLLGNGKY HSDWISNANK
MalT R    ---------- ---------- ---------- ---------- ---------- ----------

                  670        680        690        700        710        720
Consensus VRVIYWQMTG DKAAAANWLR HTAKPEFANN HFLQGQWRNI ARAQILLGEF EPAEIVLEEL
MalT WT   VRVIYWQMTG DKAAAANWLR HTAKPEFANN HFLQGQWRNI ARAQILLGEF EPAEIVLEEL
MalT R    ---------- ---------- ---------- ---------- ---------- ----------
```

```
                730        740        750        760        770        780
Consensus NENARSLRLM SDLNRNLLLL NQLYWQAGRK SDAQRVLLDA LKLANRTGFI SHFVIEGEAM
MalT WT   NENARSLRLM SDLNRNLLLL NQLYWQAGRK SDAQRVLLDA LKLANRTGFI SHFVIEGEAM
MalT R    ---------- ---------- ---------- ---------- ---------- ----------

                790        800        810        820        830        840
Consensus AQQLRQLIQL NTLPELEQHR AQRILREINQ HHRHKFAHFD ENFVERLLNH PEVPELIRTS
MalT WT   AQQLRQLIQL NTLPELEQHR AQRILREINQ HHRHKFAHFD ENFVERLLNH PEVPELIRTS
MalT R    ---------- ---------- ---------- ---------- ---------- ----------

                850        860        870        880        890        900
Consensus PLTQREWQVL GLIYSGYSNE QIAGELEVAA TTIKTHIRNL YQKLGVAHRQ DAVQHAQQLL
MalT WT   PLTQREWQVL GLIYSGYSNE QIAGELEVAA TTIKTHIRNL YQKLGVAHRQ DAVQHAQQLL
MalT R    ---------- ---------- ---------- ---------- ---------- ----------


Consensus KMMGYGV
MalT WT   KMMGYGV
MalT R    -------
```

**Fig S4. Analysis of the amino acid sequence of the MalT receptor of phage λ in wild type and resistant isolates.** Shown is a Clustal Omega 1.2.3 analysis of the amino acid sequences of the consensus (top), wild type- sensitive (middle), and resistant (bottom) *E. coli* isolates. Presented in green are the identical sequence in both isolates, in blue is the insertion sequence that leads to a frameshift in the resistant isolate, and presented in red is the site of an early stop codon in the resistant isolate.

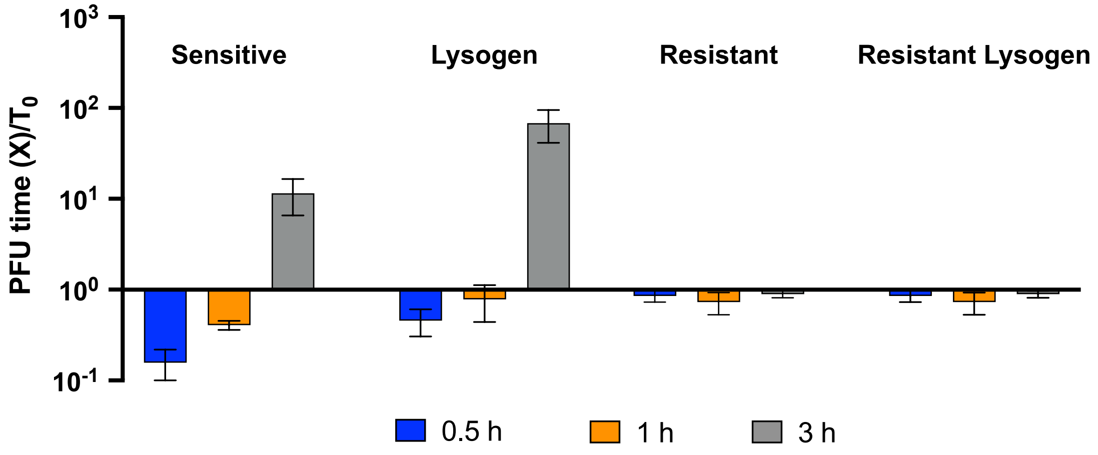


**Fig. S5. Evidence that the resistant lysogen population in Figure 2 are refractory to λ.** Ratios of the free phage density at times 0.5 (blue), 1 (orange), and 3 (grey) hours relative to that at time 0 ($T_0$). Presented are means and standard deviations of the ratios of three replicas. Infections with a low multiplicity of infection of phage λ to three known bacterial states (λ sensitive, λ lysogens, and resistant non-lysogens) as well as the presumptive resistant lysogens.

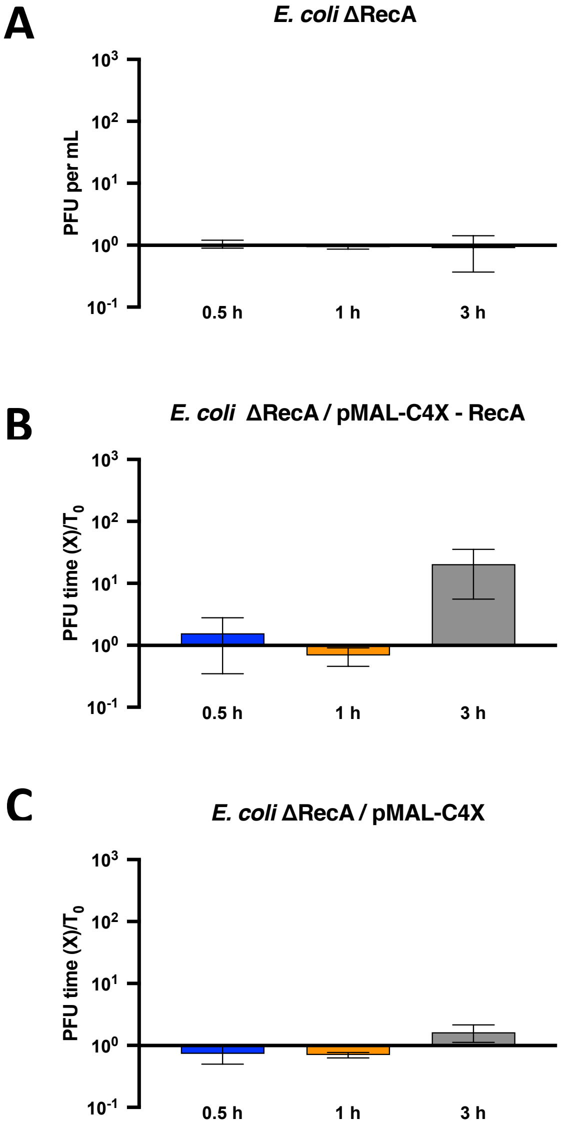


**Fig. S6. *recA* knockout and complementation assay.** Ratios of the free phage density at times 0.5 (blue), 1 (orange), and 3 (grey) hours relative to that at time 0 ($T_0$). Presented are means and standard deviations of the ratios of three replicas. Infections with a low multiplicity of infection of phage λ. **(A)** Infections of a Δ*recA E. coli* lysogenized with phage λ. **(B)** Infections of a Δ*recA E. coli* lysogenized with phage l and transformed with plasmid pMal-c4X-RecA that codes the *recA* gene in *trans*. **(C)** Infections of a Δ*recA E. coli* lysogenized with phage l and transformed with the empty plasmid pMal-c4X.

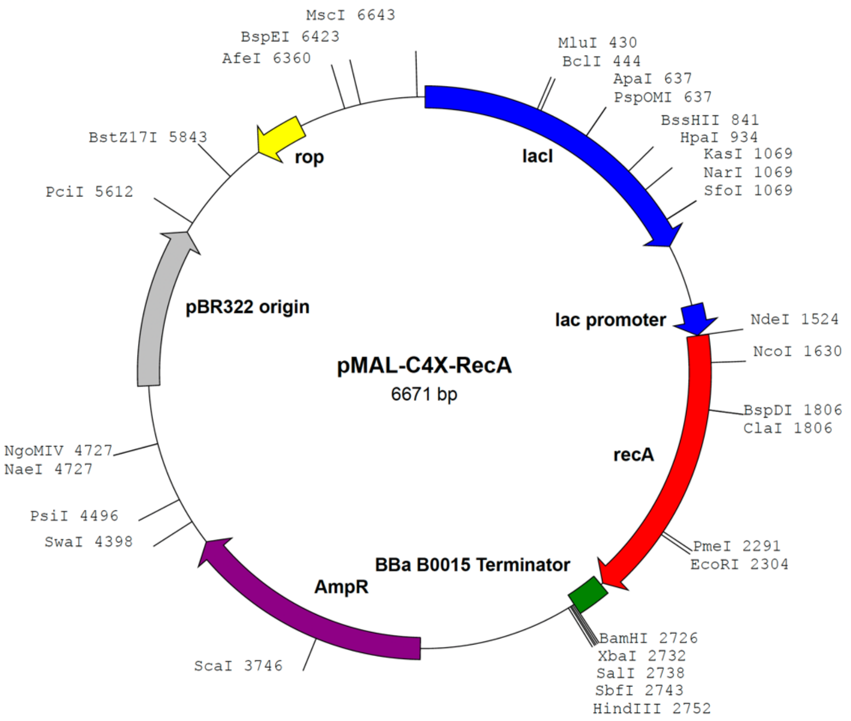


**Fig. S7. Plasmid pMal-c4X – RecA map.** Annotated diagram of the plasmid used for figure S5 to complement *recA* in *trans*.

**Supplemental Table S1. The definitions and values of the parameters used in the model**

| Parameter | Values (dimensions) | Description | Source |
|---|---|---|---|
| $v, v_{nr}, v_l, v_{lr}$ | $0.7(h^{-1})$ | Maximum growth rates | This paper |
| $\mu_{nr}, \mu_{rn}$ | $5e^{-6}, 5e^{-5}(h^{-1})$ | Transitions $N \rightarrow N_R$, $N_R \rightarrow N$ | Chaudhry *et al.* (2018) |
| $\mu_{lr}, \mu_{rl}$ | $1e^{-5}, 1e^{-5}(h^{-1})$ | Transitions $L \rightarrow L_R$, $L_R \rightarrow L$ | This paper* |
| $\delta_p, \delta_v$ | $1e^{-7}, 2e^{-7}(h^{-1} \cdot mL^{-1})$ | Adsorption rate constants | Chaudhry *et al.* (2018) |
| $\beta_p, \beta_v$ | $60\ (PFU \cdot CFU^{-1})$ | Burst sizes | This paper |
| $K$ | $1\ (\mu g)$ | Monod constant | Stewart and Levin (1973) |
| $\gamma$ | $1e^{-4}(h^{-1})$ | Induction rate | This paper* |
| $\lambda$ | $1e^{-2}(h^{-1})$ | Probability of lysogeny | This paper* |
| $e$ | $5e^{-7}(\mu g \cdot CFU^{-1})$ | Substrate conversion efficiency | Stewart and Levin (1973) |

* Estimated via numerical analysis of the results and predictions illustrated in Figure S1 and Figure S2.

**Supplemental Table S2. Wild phage genomes, their sources, and primers used to detect them**

| Wild Lysogen | Prophage(s) | Isolated Lysogen | Phage genome* | C Lysogen | Primers (5′ − 3′) |
|---|---|---|---|---|---|
| 4C10[1] | Lambda_4C10 | 1W | OM475428<br>LR595861 | 1L | Fw: CGCACGAAGAGCAGCATTAC<br>Rv: GCGTGTTATACGCCCGTTTC |
| 4A7[1] | Lambda_4A7<br>P2_4A7 | 3W | OM475430[3]<br>LR595864 | 3L | Fw: ATGGTTGGCAGTAGGCTTCC<br>Rv: CACAATGAGTGCGGCAACAA |
| 4C7[1] | Lambda | 4W | OM475432 | 4L | Fw: GACGGCTGGTATCAGGTACG<br>Rv: CTGTTCAGCAGCACGCTTTT |
| E26[2] | Lambda | E26W | OM475435 | E26L | Fw: AGCCTGTAGCTCCCTGATGA<br>Rv: GCTTATGCTTGCCGAGATGG |
| D2[2] | Lambda-like | D2W | OM475427 | D2L | Fw: CCCTTCACTGGTGACCATCC<br>Rv: GTAATGCTGCTCTTCGTGCG |
| 2H4[1] | P2_2H4 | 9W | OM475431<br>LR595869 | 9L | Fw: ACCCTAGCCTTACCCCGATT<br>Rv: CAACGGTGACCGGTTTTGAC |
| 4C9a[1] | P2_4C9 | 10W | OM475433<br>LR595883 | 10L | Fw: CGATGCGTTTCTGGCTGATG<br>Rv: CAATATTGTGCCGGTCAGCG |
| 4C9b[1] | P2-like | 11W | OM475429 | 11L | Fw: CAATATTGTGCCGGTCAGCG<br>Rv: CGATGCGTTTCTGGCTGATG |
| 4D9[1] | P2_4E6b | 12W | OM475434<br>LR595885 | 12L | Fw: TTGTCACAGGACAGACTCGC<br>Rv: CTGCAAAAACAGCGACGTGA |
| D8[2] | P2_4E6b | D8W | OM475426 | D8L | Fw: CACACAGGGACGCACTTTTG<br>Rv: TTCAGCTTTGTTGCTGTCGC |

[1] Obtained from human microbiomes.
[2] Obtained from sewage water.
[3] Only the lambda prophage was used for the C lysogen construction.
* Previous reported genomes and the generated in this study are presented in some cases.
